# Smoking and the risk for bipolar disorder: causal evidence from a bidirectional Mendelian randomization study

**DOI:** 10.1101/522268

**Authors:** Jentien M Vermeulen, Robyn E Wootton, Jorien L Treur, Hannah M Sallis, Hannah J Jones, Stanley Zammit, Wim van den Brink, Guy M Goodwin, Lieuwe de Haan, Marcus R. Munafò

**Author notes:** **Corresponding author**: Jentien Vermeulen, M.D. Amsterdam UMC Meibergdreef 9, 1105 AZ, Amsterdam, The Netherlands.

## Abstract

There is increasing evidence that smoking is a risk factor for severe mental illness, including bipolar disorder. Conversely, patients with bipolar disorder might smoke more (often) as a result of the psychiatric disorder. We aimed to investigate the direction and causal nature of the relationship between smoking and bipolar disorder we conducted a bidirectional Mendelian randomization (MR) study. Publicly available summary statistics from genome-wide association studies on bipolar disorder, smoking initiation, smoking heaviness, smoking cessation and lifetime smoking (i.e., a compound measure of heaviness, duration and cessation). We applied multiple analytical methods with different, orthogonal assumptions to triangulate results, including inverse-variance weighted (IVW), MR-Egger or Egger SIMEX, weighted median, weighted mode, and Steiger filtered analyses. Across different methods of MR, consistent evidence was found for a positive effect of smoking on the odds of bipolar disorder (smoking initiation OR_IVW_=1.46, 95% CI=1.28-1.66, P=1.44×10^-8^, lifetime smoking OR_IVW_=1.72, 95% CI=1.29-2.28, P=1.8×10^-4^). The MR analyses of the liability of bipolar disorder on smoking provided no clear evidence of a strong causal effect (smoking heaviness beta_IVW_=0.028, 95% CI= 0.003-0.053, P=2.9×10^-2^). These findings suggest that smoking initiation and lifetime smoking are likely to be a causal risk factor for developing bipolar disorder. We found some evidence that liability to bipolar disorder increased smoking heaviness. Given that smoking is a modifiable risk factor, these findings further support investment into smoking prevention and treatment in order to reduce mental health problems in future generations.

## Introduction

There is an increasing disparity in the prevalence of cigarette smoking between the general population and patients with severe mental illness, such as bipolar disorder.^1, 2^ This is important since smoking in bipolar disorder patients is associated with negative physical and mental health outcomes compared to non-smoking bipolar disorder patients.^3, 4^ Smoking has been suggested as a risk factor for developing severe mental illness, including psychosis^5^ and - to a lesser extent - bipolar disorder.^6–9^ There is also an alternative explanation, namely that patients who suffer from bipolar disorder initiate smoking or increase existing smoking. Since smoking is the most important modifiable risk factor that affects population health, more knowledge about the causal nature of the relationship between smoking and bipolar disorder could inform prevention or intervention programs.

Evidence has been found suggesting that smoking is causally related to schizophrenia and depression.^10^ In the same study, an increased liability to both depression and schizophrenia was associated with an increased risk of lifetime smoking, although evidence for this causal direction was weaker.^10^ Notably, genome-wide association studies (GWASs) have identified genetic variants coding for nicotinic acetylcholinergic receptor subunits to be associated with both schizophrenia^11^ and heavy smoking^12^ but not with bipolar disorder^13^. While a strong genetic correlation has been observed between schizophrenia and bipolar disorder^14, 15^, a genetic correlation between smoking and bipolar disorder has not been reported to date.

So far, mainly observational studies have been used to investigate the relationship between smoking and bipolar disorder.^6^ Observational results can be limited in their ability to determine causality, particularly due to residual confounding and reverse causation. Mendelian randomization (MR) is a method that aims to overcome these limitations. To the best of our knowledge, there have been no previous studies conducted into the causal relationship between smoking and bipolar disorder using this technique.

Therefore, our aim was to investigate the direction and causal nature of the relationship between smoking and bipolar disorder using a bidirectional MR approach. Based on the current evidence^6, 7, 9^ and a strong genetic correlation between bipolar disorder and schizophrenia^14^, we hypothesized that smoking is a causal risk factor for bipolar disorder. Based on conflicting clinical findings^8^ and the fact that smoking initiation often precedes the onset of bipolar disorder symptoms, we hypothesized bipolar disorder is not a causal risk factor for smoking.

## Methods

### Mendelian randomization

In this study, genetic variants are used as instrumental variables and plausibility of the findings depends on assumptions. The three key assumptions are: The multiple genetic variants associate with the risk factor of interest, there are no unmeasured confounders of the associations between genetic variants and outcome, and the genetic variants affect the outcome only through their effect on the risk factor.^16^ Pleiotropic effects refer to the effects of a genetic variant on multiple biological pathways, affecting the outcome either via the pathway under investigation, known as horizontal pleiotropy, or affect other traits through the risk factor, known as vertical pleiotropy.^27^ Horizontal pleiotropy is problematic as it violates the third key assumption of MR.

### Data sources and genetic instruments

In this MR study, multiple genetic variants that collectively explain some of the variance in the risk factors are used to investigate support for causal inferences with clinical outcomes. For smoking and bipolar disorder, genome-wide significant SNPs (P<5×10^−8^) from the most recently available GWASs were used as genetic instruments. No overlap existed in GWAS samples used for the exposure and outcome instruments.

### Smoking behavior

#### Smoking initiation

For smoking initiation, we used summary data from the GSCAN GWAS as an exposure and outcome variable. This GWAS identified 378 independent significant loci from a sample of 1,232,091 individuals of mixed ancestry: these loci explained 2.3% of the variance in smoking initiation.^17^ Data from the 23andMe sub-sample (n= 599,289) were excluded when smoking initiation was the outcome because genome-wide summary statistics were not available for the 23andMe sub-sample. Smoking initiation (yes/no) was defined as having smoked >100 cigarettes over the course of your life, smoked every day for at least a month or ever smoked regularly. The GSCAN GWAS provided effect sizes calculated from z-values weighted by the average prevalence across all studies. The effect sizes for smoking initiation as the outcome are, therefore, presented as betas and correspond to the increase of one standard deviation of prevalence of smoking initiation.^17^

#### Smoking heaviness and cessation

In order to assess causal effects of smoking heaviness and smoking cessation on the risk for bipolar disorder, the analyses would have to be stratified into groups of smokers and non-smokers. However, in the current two-sample MR study this was not possible due to a lack of data on smoking in the bipolar GWAS. Therefore, summary data about smoking heaviness and cessation from the GSCAN GWAS were used as an outcome variable only (Supplement).^17^

#### Lifetime smoking

The recently developed lifetime smoking index is a measure that applies to all individuals (i.e., ever and never smokers), and among smokers it captures smoking severity, length of exposure and cessation. This instrument does not require stratification and is therefore the best proxy to investigate a ‘dose-response’-effect on the risk for bipolar disorder. Therefore, we used lifetime smoking summary data from the GWAS in the UK Biobank.^10^ This GWAS of lifetime smoking identified 126 independent genome-wide significant SNPs in a sample of 462,690 individuals with European ancestry.^10^ It has been validated in an independent sample, where the 126 SNPs explained 0.36% of the variance in lifetime smoking, and against positive control disease outcomes (e.g., lung cancer).^10^

### Bipolar disorder

For bipolar disorder as an exposure, we used summary statistics from the most recent GWAS comprising 20,352 cases from Europe, North-America and Australia and 31,358 controls of European descent.^13^ The GWAS found 30 independent, genome-wide significant loci in the combined metaanalysis of the discovery GWAS and follow-up samples. The polygenic risk score, including all measured genetic variants, accounted for about 8% of the variance in diagnosis. When bipolar disorder was the outcome, we used the one publicly available GWAS summary data made available by Psychiatric Genomics Consortium with a similar number of bipolar disorder cases (20,129 bipolar disorder cases and 21,524 controls).^14^

### Statistical Analyses

We used the MR Base Package^18^ in R version 3.5.1^19^ to harmonise SNP information and conduct bidirectional summary level MR analyses between smoking and bipolar disorder. The magnitude of the causal effect can be estimated by dividing the genetic variant-outcome association by the genetic variant-risk factor association.^20^ Results were compared across the following methods: inverse-variance weighted (IVW), MR Egger^21^ or Egger SIMEX^22^, weighted median^23^, weighted mode^24^ and MR RAPS. A consistent direction of effect across all of these methods strengthens causal evidence as each method makes different assumptions about pleiotropy.^26^ Cochran’s Q (for IVW) and Rucker’s Q (for MR Egger) statistics were therefore calculated to investigate heterogeneity between SNP causal effects and thus potential evidence of horizontal pleiotropy. Evidence of directional pleiotropy (whereby pleiotropic effects are biasing the estimates in the same direction) was also assessed using the MR Egger intercept. The difference between Cochran’s Q and Rucker’s Q statistics (Q’) was used to assess the extent to which MR Egger was a better fit to the data than the IVW method. The validity of the MR Egger method was also evaluated using the weighted and unweighted regression dilution I^2^_GX_ statistic.^28^ For values of I^2^_GX_ < 0.9, MR Egger SIMEX corrections are presented using the highest of the two regression dilution statistics. For values of I^2^_GX_ < 0.6, MR Egger and MR Egger SIMEX are unreliable and so neither are performed. Furthermore, mean F-statistics were calculated to assess the strength of the genetic instruments used.^29^ As a rule of thumb, values of the mean F-statistic above 10 indicate that the results do not suffer from substantially weak instrument bias.^20^ Lastly, we applied Steiger filtering to remove the genetic variants that explain more variance in the outcome than the exposure to avoid possible reverse causation.^30^

## Results

### Summary level Mendelian randomization of smoking on bipolar disorder

Across different methods of MR analyses, consistent evidence was found for a positive effect of smoking initiation on the odds of bipolar disorder (OR_IVW_=1.46, 95% CI=1.28-1.66, P=1.4×10^-08^) (see Table 1 and Supplemental Figures S1). MR analyses using lifetime smoking as the exposure variable showed significant effects in the same direction with a similar effect size (OR_IVW_=1.72, 95% CI= 1.89-2.28, P=1.8×10^-04^) (see Table 1 and Supplemental figure S7). The I^2^_GX_ suggested to use MR Egger SIMEX results (see Supplemental Table S1 and Table 1). Tests for heterogeneity suggest evidence for significant horizontal pleiotropy (see Supplemental Table S1), but MR Egger intercepts suggest that bias of the estimates by pleiotropy is not likely (see Supplemental Table S2). To explore whether the genetic variants for lifetime smoking and smoking initiation explained more variance in the exposure than in the outcome, Steiger filtering was applied. For smoking initiation, 20 SNPS dropped out and 342/362 SNPS (94%) could be included after Steiger filtering. For lifetime smoking, 12 SNPS dropped out and 107/119 SNPs (90%) could be included in the analyses following Steiger filtering. The analyses were repeated across methods and similar effects were observed for smoking initiation and lifetime smoking on bipolar disorder (smoking initiation OR_IVW_=1.44, 95% CI=1.29-1.62, 1.0×10^-04^; lifetime smoking OR_IVW_=1.47, 95% CI=1.17-1.83, P =7.9×10^-04^) (see Supplemental Table S5). Again, MR Egger was not conducted as I^2^_GX_ values were below 0.9 (see Supplemental Table S1).

**Table 1.**
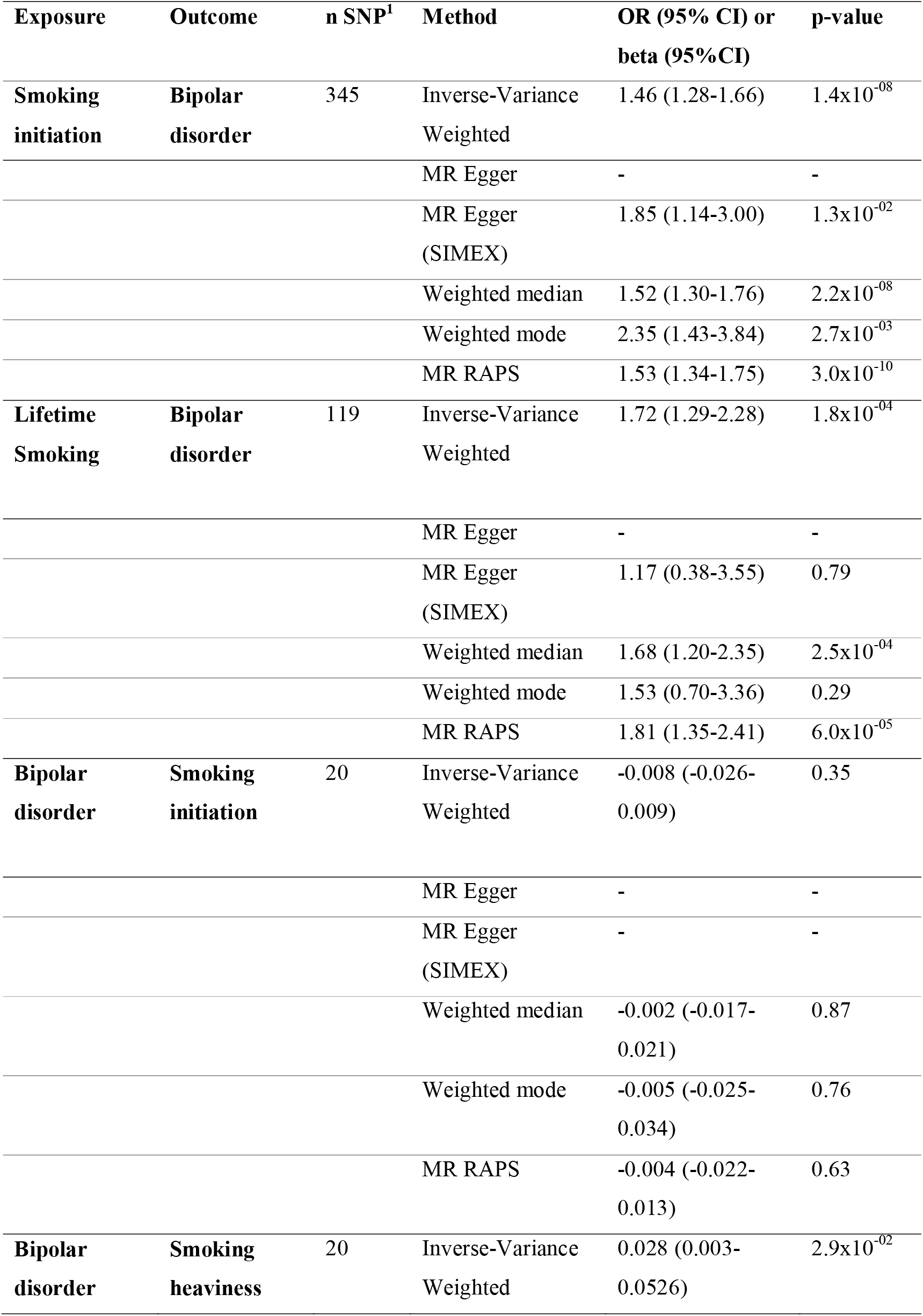
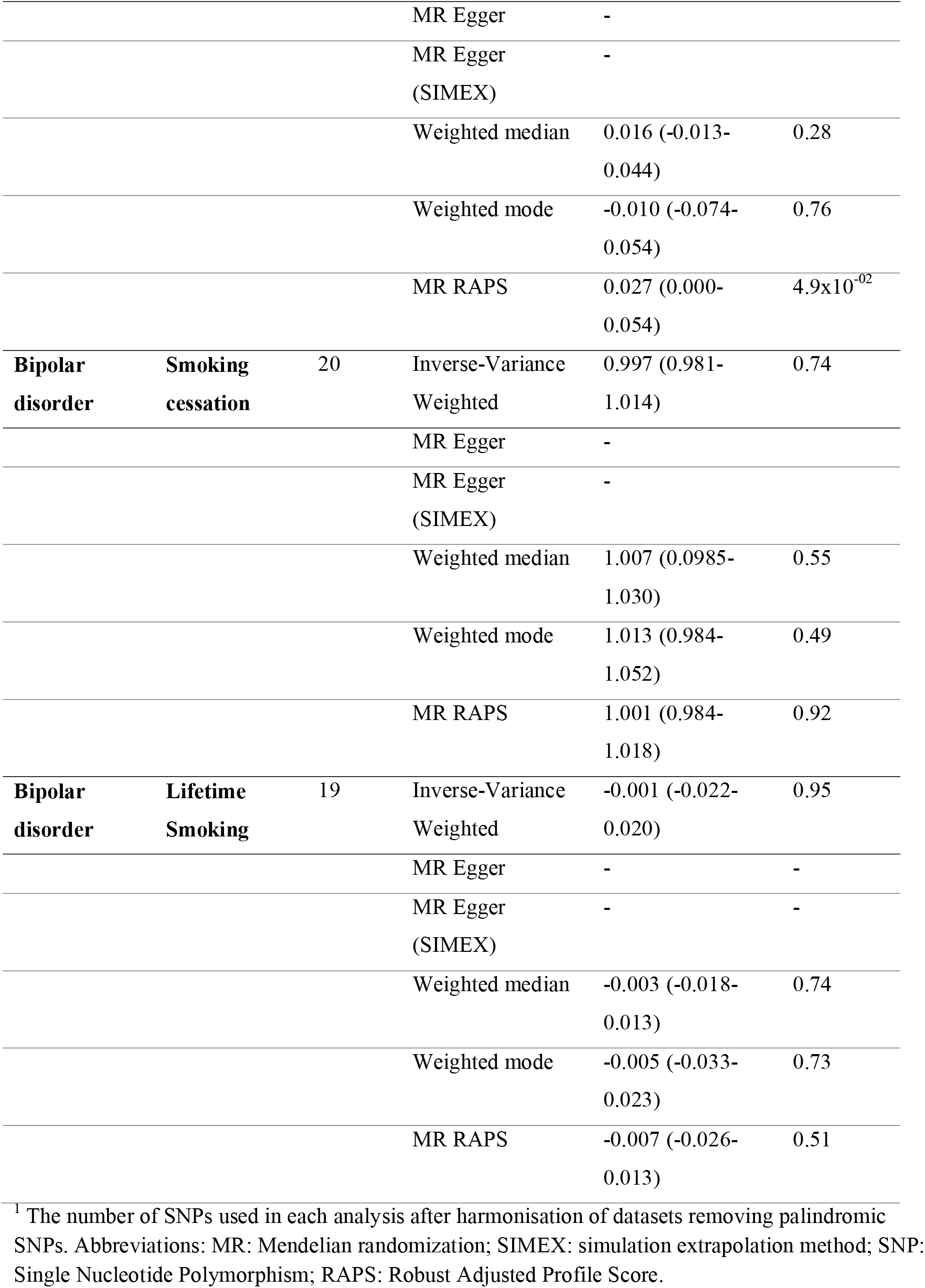
Bidirectional Mendelian randomization analyses of the effects of smoking and bipolar disorder using summary level data.

### Summary level Mendelian randomization of bipolar disorder on smoking

The MR analyses of liability of bipolar disorder on risk of smoking initiation (beta_IVW_= −0.008, 95% CI= −0.026-0.009, P=0.35), lifetime smoking (beta_IVW_= −0.001, 95% CI= −0.032-0.030, P=0.95) and smoking cessation (OR_VW_= −0.003, 95% CI= 0.981-1.014, P=0.74) revealed no evidence of a causal effect (see Table 1 and Supplemental Figures S4 and S10). MR analyses of liability of bipolar disorder on smoking heaviness (beta_IVW_= 0.028, 95% CI= 0.003-0.053, P=2.9×10^-2^) showed significant, small effects for MR IVW and MR RAPS but not MR weighted median or MR weighted mode (Table 1 and Supplemental Figure S13). The I^2^_GX_ suggested not to use MR Egger or MR Egger SIMEX results (see Supplemental Table S1). The tests for heterogeneity indicated no evidence for significant heterogeneity (Supplemental Table S2) and MR Egger intercepts suggested that bias of the estimates by horizontal pleiotropy is not likely (Supplemental Table S3). Steiger filtering indicated that all genetic variants for bipolar disorder explained more variance in bipolar disorder than smoking outcomes. The mean F-statistic also suggested that results were not subject to weak instrument bias (Supplemental Table S4). Although we did not find an effect using multiple genome-wide significant genetic variants, we noted that one SNP (rs3804640) stood out in particular showing a large, positive effect in the single SNP analysis of bipolar disorder to smoking initiation and lifetime smoking (Supplemental Figure S5 and S11). This SNP is an intron in the gene *CD47*, which encodes a membrane protein shown to play a role in synaptic plasticity in the hippocampus.^18–20^

## Discussion

The current study shows that smoking is a causal risk factor for bipolar disorder by applying MR to two measures of smoking behaviour using GWAS summary statistics. In contrast, liability to bipolar disorder does not have a causal effect on smoking initiation, smoking cessation or lifetime smoking. We found some evidence that suggested a small effect of liability for bipolar disorder on increased smoking heaviness. To the best of our knowledge, this is the first study providing robust evidence on the direction of the association between smoking and bipolar disorder.

### Mendelian randomization of smoking on bipolar disorder

When comparing the current findings to those of a previous study evaluating the relation between smoking and other severe mental illnesses, the effect sizes we found are comparable to those testing lifetime smoking and smoking initiation as a risk factor for schizophrenia and major depressive disorder.^10^ The evidence for smoking as a risk factor for bipolar disorder has been meager.^7–9^ However, one conventional population-based birth cohort study found that maternal smoking was a risk factor for bipolar disorder in the offspring.^6^ To rule out the effect of maternal smoking as an explanation of our results, future cohorts with measures of maternal and offspring smoking with enough bipolar cases are needed. In line with our findings, a study that used polygenic risk scores of smoking to predict bipolar disorder revealed a significant association but not vice versa.^21^ Although this is suggestive of a causal effect, the current study adds to the body of evidence by overcoming residual confounding and providing an estimate of the causal effect size.

### Mendelian randomization of bipolar disorder on smoking

Although other studies have found an effect of bipolar disorder on smoking behavior^8^, the current study did not replicate this finding for most smoking behaviours. The lack of an effect indicates that liability to bipolar disorder is not a risk factor for smoking initiation, cessation or lifetime smoking. From a methodological point of view, one might explain the lack of a clear effect of the liability to bipolar disorder on smoking initiation, cessation and lifetime smoking by using weaker instruments for bipolar disorder as compared to smoking. This would result in reduced statistical power to detect a causal effect. However, we argue that this explanation is unlikely since the effect sizes consistently show values around the null, confidence intervals were small and the mean F-statistic, which is a proxy for the strength of the instrumental variable, showed an acceptable value. The results of the liability to bipolar disorder on smoking heaviness suggested from two MR methods a significant, small effect and from two other methods no significant effect. This could indicate that liability to bipolar disorder increases smoking heaviness or there might be a bias from pleiotropy. Of note, MR typically investigates long-term effects of liability to bipolar disorder on life-time smoking behaviour and firm conclusions on acute transient effects (e.g., more severe bipolar symptoms increasing heaviness of smoking) could, therefore, not be drawn.

### Strengths and limitations

A strength of this study is the substantial power, since we used summary data from recent, large GWAS consortia to conduct a summary level MR.^22^ Also, using genetic variants to determine the effects reduces the chance of reverse causation and residual confounding.^23^ Furthermore, a consistent effect across different methods increases the confidence in most of our findings.^24^ Some limitations should be noted. First, the GWAS were conducted in individuals from European ancestry which could limit generalizability to other populations. Second, although the smoking instruments capture smoking initiation, duration, heaviness and cessation, the instruments cannot be used to determine effects at specific life stages. The results should be interpreted as cumulative effects of smoking over the life course (potentially from conception and potentially from second hand smoke from family members). In other words, it is not certain that our findings regarding smoking are confined to a specific period. Third, we assumed that the exclusion restriction criteria hold since the MR-Egger intercept test for directional horizontal pleiotropy was not significant. Still, pleiotropy could be in place as regression dilution values pointed that Egger results might not be trustworthy. The underlying biological process of mental illnesses are relatively poorly understood which complicates the possibility to disentangle the plausibility of potential pathways. For example, genetic variants that were significantly associated with smoking initiation coded for several biologically plausible pathways such as acetylcholine nicotinic receptors and glutamate pathways that affect reward processing and addiction.^17^ However, general biological pathways such as sodium, potassium and calcium voltage-gated channels were also implied, potentially overlapping with horizontal pleiotropic pathways. Fourth, genetic vulnerability for smoking could also be related to environmental factors that are associated with the outcome which could lead to spurious causal effects. Nevertheless, the consistent findings across all sensitivity analyses with different assumptions point towards robust findings. Lastly, sensitivity analyses into bipolar subtypes could not be performed due to a low number of associated genetic variants. Future GWAS with larger sample sizes are required to potentially overcome this issue.

### Possible mechanisms

There are several, non-mutually exclusive, plausible explanations for causal pathways from smoking to bipolar disorder.^7–9^ First, nicotine binds to subtypes of the nicotinic acetylcholinergic receptor which is implicated in the regulation of central nervous system pathways relevant to mood disorders.^25^ Although the quality of the evidence is low, several case studies have related the use of a partial nicotine receptor agonist (varenicline) to induced mania or new onset of bipolar disorder.^26–31^ Second, nicotine exposure leads to desensitization of the nicotine acetylcholinergic receptor by phospohorylation of subunits.^32^ These dysmorphic receptors then have reduced binding capacity for endogeneous acetylcholine: craving on drug withdrawal is clear evidence of changed function in reward pathways. The reduced functioning of these receptors could have an effect on neurotransmitter levels and subsequently lead to a higher vulnerability to develop bipolar disorder, although evidence about this topic is lacking. Third, burning tobacco releases toxic compounds which induce inflammation and oxidative stress which may be associated with bipolar disorder.^25^

### Clinical implications

Many hypotheses exist to explain the high prevalence of smoking in severe mental illness.^33^ The lack of a consistent, bidirectional effect and the causal evidence that smoking initiation liability increases the risk for bipolar disorder point toward the two-hit or diathesis-stress model.^34^ This implies a neurobiological vulnerability for severe mental illness^35^ interacting with an environmental stressor, such as smoking, leading to an increased vulnerability for severe mental illness.^34^ Several studies have shown that smoking in adolescents and adults with bipolar disorder is associated with earlier age at onset of mood disorder, greater severity of symptoms^36–41^, poorer functioning, comorbid substance abuse^42^, suicide attempts^43^ and other adverse outcomes such as readmission^44^. Our results underline the importance of strategies to prevent adolescents from starting smoking and treat smoking in bipolar patients. The frequently mentioned reason for smoking by bipolar patients is stress-coping behavior^45^ and a tailored prevention strategy might be helpful in learning healthy coping strategies. Regardless of the mechanism which may eventually be confirmed, our findings underline the importance of prevention and treatment programs to reduce smoking and its consequences. Investing in a smoke free future could reduce mental health problems in future generations.

